# Tetratricopeptide repeat domain 36 protects renal tubular cells from cisplatin-induced apoptosis via maintaining mitochondrial homeostasis

**DOI:** 10.1101/2021.04.25.441359

**Authors:** Xin Yan, Rui Peng, Dayu Tian, Lei Chen, Qingling He, Qianyin Li, Qin Zhou

## Abstract

The apoptosis of proximal tubule epithelial cells (PTECs) is a critical event of acute kidney injury (AKI). Tetratricopeptide repeat domain 36 (TTC36) with three tetratricopeptide repeats is evolutionarily conserved across mammals, which functions as a chaperone for heat shock protein 70. We have revealed that TTC36 is specifically expressed in PTECs in our previous work. There are few studies about the role TTC36 played in AKI. Therefore, in this study, we investigated the function of TTC36 in the apoptosis of HK2 cells, which are derived from the human proximal tubule. Firstly, we observed that TTC36 was obviously down-regulated and was negatively related to the kidney damage degree in a mouse model of acute kidney injury established by ischemia/reperfusion. In addition, TTC36 overexpression protected HK2 cells against cisplatin-induced apoptosis. Moreover, we discovered the mechanism that TTC36 mitigated cisplatin-triggered mitochondrial disorder via sustaining the membrane potential of mitochondria and mitochondrial autophagy-related gene expression. Collectively, these results suggested that TTC36 plays a protective role in the cisplatin-induced apoptosis of renal tubular cells through maintaining the mitochondrial potential and mitochondrial autophagy-related gene expression. These observations highlight the essential role of TTC36 in regulating PTEC apoptosis and imply TTC36/mitochondrial homeostasis axis as a potential target for the therapeutic intervention in AKI.

## Introduction

Acute kidney injury (AKI), with the characteristic of the rapid decline of glomerular filtration rate (GRF), is a worldwide clinical syndrome accompanied by the sudden increase of serum creatinine (SCr) and blood urea (BUN)(1,2). AKI, primarily caused by ischemia, sepsis, and nephrotoxicity, not only leads to approximately 1.7 million deaths every year but also makes patients prone to chronic kidney disease (CKD)(1,3,4). However, there is still no efficient remedy for curing AKI. Therefore, it is necessary to investigate the underlying mechanisms of AKI and excavate the potential strategy for the prevention and treatment of AKI.

During the progression of AKI, the dysfunction and apoptosis of proximal tubular epithelial cells, which are intensively packed with mitochondria, are regarded as a key event(5). Synthesizing adenosine 5’-triphosphate (ATP) via electron transport and oxidative phosphorylation (OXPHOS) in concert with the oxidation of metabolites by the tricarboxylic acid cycle and catabolism of fatty acids by β-oxidation, mitochondria play a critical role in supporting cellular energy-intensive processes, including generating ion gradients and reabsorbing ions(6–8). Adjusted by mitophagy, autophagy-related clearance of impaired mitochondria, mitochondrial biogenesis and mitochondrial dynamics, mitochondrial homeostasis are vital for cellular homeostasis and function(9). However, the disorder of mitochondrial homeostasis and the damage of PTECs’ integrity could contribute to energy metabolism falling apart, which finally facilitates or even leads to AKI(10–12). Given the significance of mitochondria in energy homeostasis, it is a potential and promising strategy that mitigating AKI via targeting mitochondrial metabolism.

Tetratricopeptide repeat domain 36 (TTC36) is a conserved protein with three tetrapeptide repeats, which serves as a chaperone of heat shock protein 70. TTC36 primarily expressed in liver and renal proximal tubular epithelial cells in kidneys(13). Current studies have observed that it participates in the metabolism of tyrosine by interacting with 4-hydroxyphenylpyruvic acid dioxygenase (HPD) and reducing the binding of serine/threonine kinase 33 (STK33) to HPD to block the degradation of HPD in hepatocyte(14). It is worth noting that TTC36 is restrictively expressed in renal proximal tubules in kidney. However, there is no research revealing the role of TTC36 in acute kidney injury.

In this study, we employed HK2 cells, which are derived from human proximal tubule, and established an animal model to uncover the role of TTC36 in AKI. We revealed that TTC36 protects HK2 cells against cisplatin-induced apoptosis via maintaining mitochondrial membrane potential and mitochondrial autophagy-related gene expression, implying a promising access to treat AKI.

## Materials and Methods

### Materials

Rabbit polyclonal antibody anti-TTC36 was made in previous work(15). Mouse anti-Bcl2 (sc-7382), anti-Bax (sc-20067), and anti-Phospho-Bcl2 (sc-293128) were purchased from Santa Cruz Biotechnology (Santa Cruz, CA, USA). Rabbit antibodies against Caspase-3 (9662S) and Caspase-9 (9502S) were obtained from Cell Signaling Technology (Danvers, MA, USA). Mouse anti-Flag (AE005) and anti-GAPDH (AC002) were obtained from Abclonal (Wuhan, China). Mouse anti-β-Actin (HC201) was obtained from TransGen Biotech (Beijing, China). JC-1 (HY-15534), Cisplatin (CP) (HY-17394) were obtained from MedChemExpress (New Jersey, USA). For Immunohistochemical (IHC) staining, SP kit (SP-9001) and Diaminobenzidine (DAB) coloring kit (ZLI-9017) were purchased from ZSGB-BIO (Beijing, China). Serum creatinine (SCr) detection Kit (C011-2-1) and blood urea nitrogen (BUN) detection kit (C013-2-1) were obtained from Nanjing Jiancheng Bioengineering Institute (Nanjing, China).

### DNA Construction

To construct the overexpression vector of *TTC36*, human *TTC36* (NM_001080441.4) was amplified with cDNA and subcloned into the CMV vector.

#### Mice and AKI model

All animals were reared under the condition of a standard laboratory where food and water were sufficiently supplied. 8-week-old male wild-type (WT) C57BL/6 mice were obtained from the Laboratory Animal Centre of Chongqing Medical University (No. SYXK2018-0003, Chongqing, China). *Ttc36* knockout (*Ttc36*^-/-^) mice were generated in previous work(16), these mice develop normally with a normal lifespan even though they will suffer from Hypertyrosinemia when 8-12 month-old. A total of 22 mice were used in this study, including 19 WT and 3 *Ttc36*^-/-^ C57BL/6 mice. To evaluate the correlation of TTC36 expression with renal function, 8-week-old male WT C57BL/6 mice were assigned to 4 groups (Sham, n = 4; 2 days after IR-treated, n = 4; 7 days after IR-treated, n=4; 14 days after IR-treated, n=4). For the isolation of primary renal tubular epithelial cells, 8-week-old male WT and *Ttc36*^-/-^ C57BL/6 mice were sacrificed (WT mice, n = 3; *Ttc36*^-/-^ mice, n = 3). All animal related experiments were approved by Animal Ethical Commission of Chongqing Medical University. The mice were anesthetized with ether and warmed by a heating pad at 37.0 °C during the surgery.

To establish the acute kidney injury, bilateral renal arteries were separated and were clamped for 30 min(17). In the sham group, the mice were treated with the same way except arteries clamping. Mice were euthanized by cervical dislocation after anesthetized with ether, then the blood were collected for the detection of SCr and BUN and kidneys were obtained for Hematoxylin-Eosin (HE) staining, immunohistochemical (IHC) staining, mRNA quantification, and western blotting.

### Cell culture and treatment

HK2 cells, which derived from the human proximal tubule, were cultured in Dulbecco’s modified Eagle’s medium-F12 (DMEM/F12) medium (Gibco) supplemented with 10% fetal bovine serum (Biological Industries) and 1% penicillin-streptomycin (HyClone). The HEK293T cells, packaging cells, were maintained in DMEM medium (Gibco) containing 10% fetal bovine serum (Biological Industries) and 1% penicillin-streptomycin (Gibco). To construct TTC36 stably overexpressed cells, HEK293T cells were transfected with TTC36 overexpression vectors and lentivirus backbone vectors using TurboFect™ Transfection Reagent (ThermoFisher Scientific). Then, for the enrichment of lentivirus, the supernatant were collected and purified by ultra-high-speed centrifugation (25,000 × g for 2h at 4°C) after transfection for 48h. HK2 cells were infected with lentivirus using 8 μg/mL polybrene (Sigma). To select the stable clones, HK2 cells were treated with puromycin (Invitrogen). To knock down the expression of TTC36, si-RNA for silencing TTC36 (si-TTC36, sequence 5’-GGAAGAACGAGAAGAAGAUGA-3’) were synthesized by GenePharma (Jiangsu, China), and transfected into HK2 cells with Lipofectamine 2000 Transfection Reagent (ThermoFisher Scientific) according to the manual. In order to establish the in vitro model of AKI, HK2 cells were treated with a series concentration of cisplatin, then we used 25μM cisplatin to treat HK2 cells for 20 hours to induce acute tubular injury.

### Isolation of Mouse Primary Renal Tubular Cells

Primary renal tubular cells were isolated from 8-week-old wild type and *Ttc36*^-/-^ C57BL/6 mice. In Brief, after starved overnight, the mice were anesthetized with ether and their kidneys was perfused with 20 mL perfusion buffer (1% penicillin-streptomycin (HyClone) in PBS) through the left ventricle and then perfused with 30 mL digestion buffer (0.13 mg/mL collagenase type II (Sigma), Hank’s balanced salt solution (HyClone) with 5 mM Ca^2+^ and 1.2 mM Mg^2+^). After perfused, the kidneys were removed from the abdominal cavity. After the renal capsules and medulla were removed, the cortex was cut into tiny pieces and incubated with digestion buffer at 37 °C for 10 min. The tubular cells were collected using filters followed by centrifuging at 50×g for 5 min. After that, cells were suspended, transferred into collagen type I coated 100 mm dishes (coating buffer, 4% collagen type I (Corning) and 0.2% glacial acetic acid in PBS), and cultured in DMEM/F12 (Gibco) medium containing 20% FBS(Gibco) and 1% penicillin-streptomycin (HyClone).

### Hematoxylin and Eosin (H&E) Staining and IHC Staining

The kidneys were harvested, fixed in 4% formaldehyde, embedded in paraffin, and cut into 4 μm sections. To process the H&E staining, have dewaxed and rehydrated, the sections were stained with hematoxylin (2 min), soaked with acid alcohol (2 sec), soaked in lithium carbonate (2 min) and 80% ethanol (1 min) in sequence, and counterstained with eosin for 1min. For IHC staining, after pre-heated at 65°C for 2h, the sections were dewaxed and rehydrated, heated at 100°C for 20min in the presence of citric acid buffer (pH = 6.0) to retrieve antigen, placed at room temperature for 4h to cool down, treated with hydrogen-peroxide-solution for 10min, coated with goat serum for blocking, incubated with the antibody at 4°C for overnight, washed with PBST(0.05% Tween20 in PBS)for three times, coated with appropriate secondary antibody labeled with biotin for 15min, incubated with streptavidin-conjugated with horseradish peroxidase (HRP) at room temperature for 15min. After that, the sections were stained using a DAB coloring kit.

### Western Blotting

Cells were lysed with lysis buffer (50 mM Tris–HCl pH 7.5, 1% SDS, 1% TritonX-100,150 mM NaCl, 1 mM dithiothreitol, 0.5 mM EDTA, 100 mM PMSF, 100 mM leupeptin, 1 mM aprotinin, 100 mM sodium orthovanadate, 100 mM sodium pyrophosphate, and 1 mM sodium fluoride) after rinsed three times with cold PBS, then the extraction were collected to 1.5 ml microtube and heated at 95 °C in mental bath (ThermoFisher Scientific) for 10 min. After that, the protein was centrifuged at 10,000×g for 10 min and quantified by BCA Protein Assay Kit (Thermo Fisher Scientific). The same quantity (about 15 μg) protein of each sample was separated with 12% sodium dodecyl sulfate-polyacrylamide gel and transferred to polyvinylidene difluoride membranes (Millipore). After blocked by 5% fat-free milk (Sangon Biotech) in Tris-buffered saline containing 0.08% Tween 20 (Sigma) at room temperature for 2h, the membranes were incubated with primary antibodies at 4°C overnight, washed three times with TBST,10min each, incubated with appropriate secondary antibodies coupled with HRP for 1 hour at room temperature, and washed with TBST again. The target bands were detected with Smart-ECL basic (Smart-Lifesciences) using Image Lab software program (Bio-Rad). Actin or GAPDH was used for the loading control. For the detection of Bcl2 and p-Bcl2, the PVDF membranes were first incubated with anti-p-Bcl2 antibody to detect it, then these PVDF membranes were washed with stripping buffer to stripe the bands and incubated with anti-Bcl2 antibodies after blocked with 5% fat-free milk. For integral optical density analysis, the bands were scanned and calculated by Image J software.

### Mitochondrial Membrane Potential Assay

The membrane potential (ΔΨm) of mitochondrial was determined by JC-1 which is a fluorescent dye and can selectively enter mitochondria. In brief, when the ΔΨm is relatively low, JC-1 forms monomers and emits green fluorescence. On the contrary, when the ΔΨm is high, it will aggregate and transmit red fluorescence. After treatment, HK2 cells were incubated with JC-1 according to the manual, and detected by a CytoFLEX flow cytometry (BECKMAN COULTER)

### Cell Viability Assay

After treated with cisplatin, the cell viability assay was performed using methyl thiazolyl tetrazolium (MTT). Briefly, cells were cultured in a 96-well plate, and each well was added 20 μl 5 mg/ml MTT, incubated at 37°C for 4 h. Then, the medium was discarded and each well was added 150 μl dimethylsulfoxide (DMSO), shaking at low speed for 10 min, and determined with MULTISKAN GO (ThermoFisher Scientific) at OD 490 nm.

### Apoptosis Assay

Cells undergoing apoptosis were quantitatively determined with Annexin V PE/7-AAD kit (Solarbio). Digested with trypsin and washed with PBS for three times, cells were incubated with the reagent according to the manufacturer’s instructions, and detected with CytoFLEX flow cytometry (BECKMAN COULTER).

### Quantitative Real-Time Polymerase Chain Reaction (RT-qPCR)

For the detection of specific gene’s mRNA expression level in HK2 cells and kidney tissue, cells or tissue were lysed with TRIzol reagent (Invitrogen). The total RNA was extracted using chloroform and isopropanol, washed with 75% ethyl alcohol, and dissolved with DNase/RNase-free water (Solarbio). To perform RT-qPCR, the total RNA was used for synthesizing cDNA with RevertAid RT Reverse Transcription Kit (Thermo Fisher scientific) under the guidance of manufacturer’s instructions. After that, the cDNA library was amplified in the presence of specific primers and 2x SYBR Green qPCR Master Mix (bimake) with a CFX96 real-time PCR detection system (Bio-Rad). The relative expression level of specific genes was analyzed relative to the mean critical threshold (CT) values of the 18S gene. Primer sequences of specific genes were listed in Supplementary Table S1.

### Statistical Analyses

All experiments were repeated independently three times. Data were exhibited as mean ± standard deviation using GraphPad Prism 8 software, and the Statistically significant differences were calculated by analysis of variance (ANOVA), followed by a Student’s *t*-test using IBM SPSS Statistics 20 software. Differences were regarded as significant with *p*< 0.05. * *p* < 0.05, ** *p* < 0.01, *** *p* < 0.001, and **** *p* < 0.0001.

## Results

### The reduction of TTC36 expression in murine renal tubular cells was related to AKI

First, we established the in vivo model of AKI with the bilateral renal artery and vein clamping for 30 minutes, as shown in Fig. 1A. Next, we detected the concentration of serum creatinine (SCr) and blood urea nitrogen (BUN) in mice at the indicated time, and an apparent increasement of SCr and BUN were observed in IR-treated mice after 2 days compared to those in the Sham group (Fig. 1, B and C). At the same time, a significant reduction of TTC36 expression was detected in kidneys from mice with IRI, and recovery of TTC36 expression along with the decrease of SCr and BUN concentration was found (Fig. 1, B to E).

**Figure 1.**
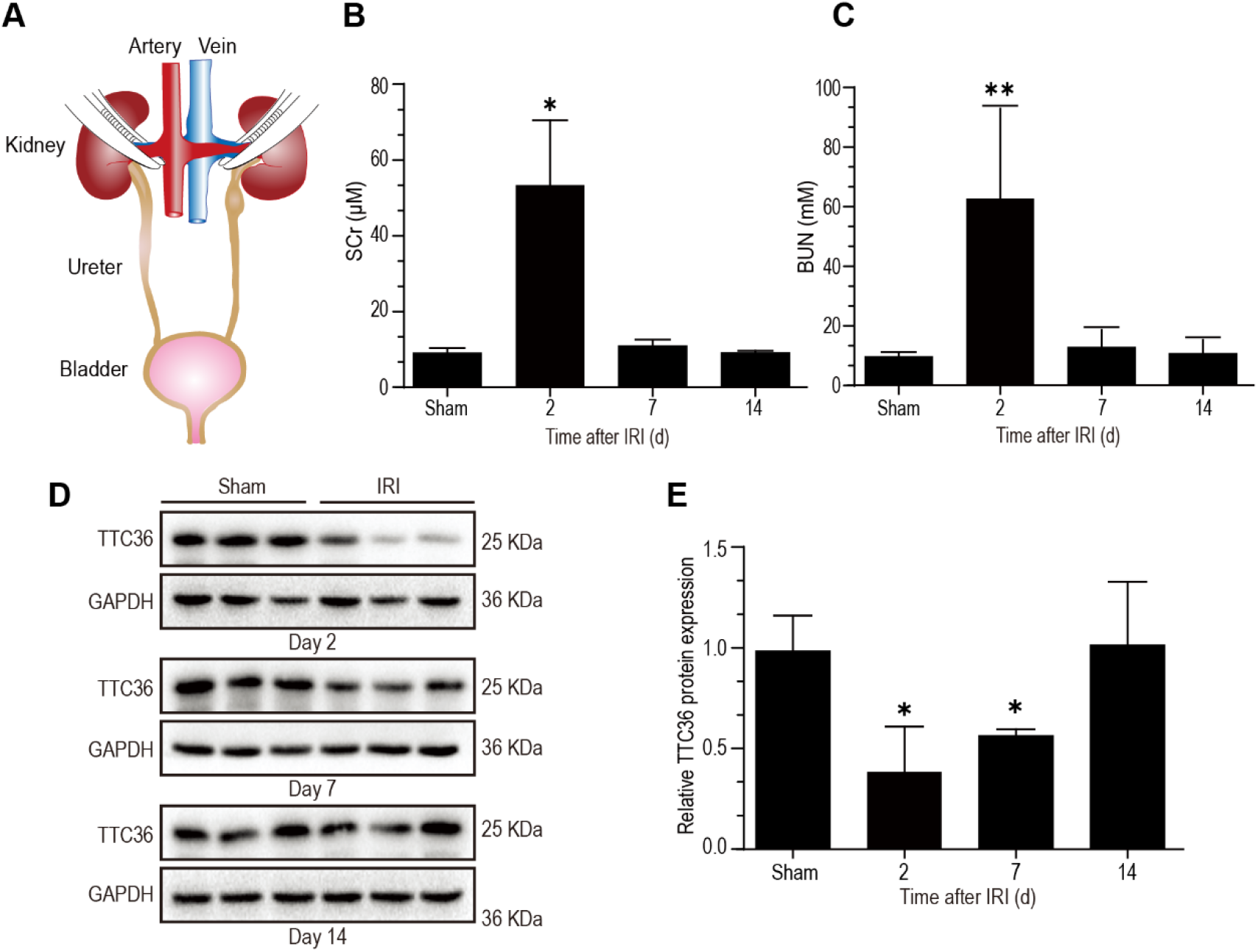
The reduction of TTC36 expression in murine renal tubular cells was related to AKI. (A) surgery strategy of IR to induce AKI. (B) the Concentration of SCr in IR-induced mice (Sham and 2, 7 and 14 days after IR-treatment; n=4 mice per group). (C) the Concentration of BUN in IR-induced mice (Sham and 2, 7 and 14 days after IR-treatment; n=4 mice per group). (D) Western blotting for TTC36 in the kidneys of IR-treated mice (Sham, 2, 7 and 14 days after IR-treatment; n = 3 mice per group). (E) Quantitative analysis of the TTC36 expression related to GAPDH through detecting the integral optical density of (D). Data are expressed as means + SD. Statistically significant differences were determined by Student’s *t*-test and one-way ANOVA. \**p* < 0.05 and \*\**p* < 0.01 versus Sham group. Results are representative of at least three independent experiments. SCr, serum creatinine; IR, ischemia-reperfusion; BUN, blood urea nitrogen; TTC36, tetratricopeptide repeat domain 36; GAPDH, glyceraldehyde-phosphate dehydrogenase; AKI, acute kidney injury; SD, standard deviation; ANOVA, analysis of variance.

### The expression of TTC36 was negatively corelated with the degree of kidney injury

In line with the above data, an obvious renal tubular injury accompanied by the downregulation of TTC36 expression was confirmed in histological staining (Fig. 2, A and B). Further analysis demonstrated that the expression of TTC36 in tubular cells was negatively correlated with the concentrations of SCr (r = −0.7224, *p* < 0.0001) and BUN (r = −0.6870, *p* < 0.001) (Fig.2 C and D), suggesting that the down-regulation of TTC36 is associated with the pathogenesis of acute tubular injury.

**Figure 2.**
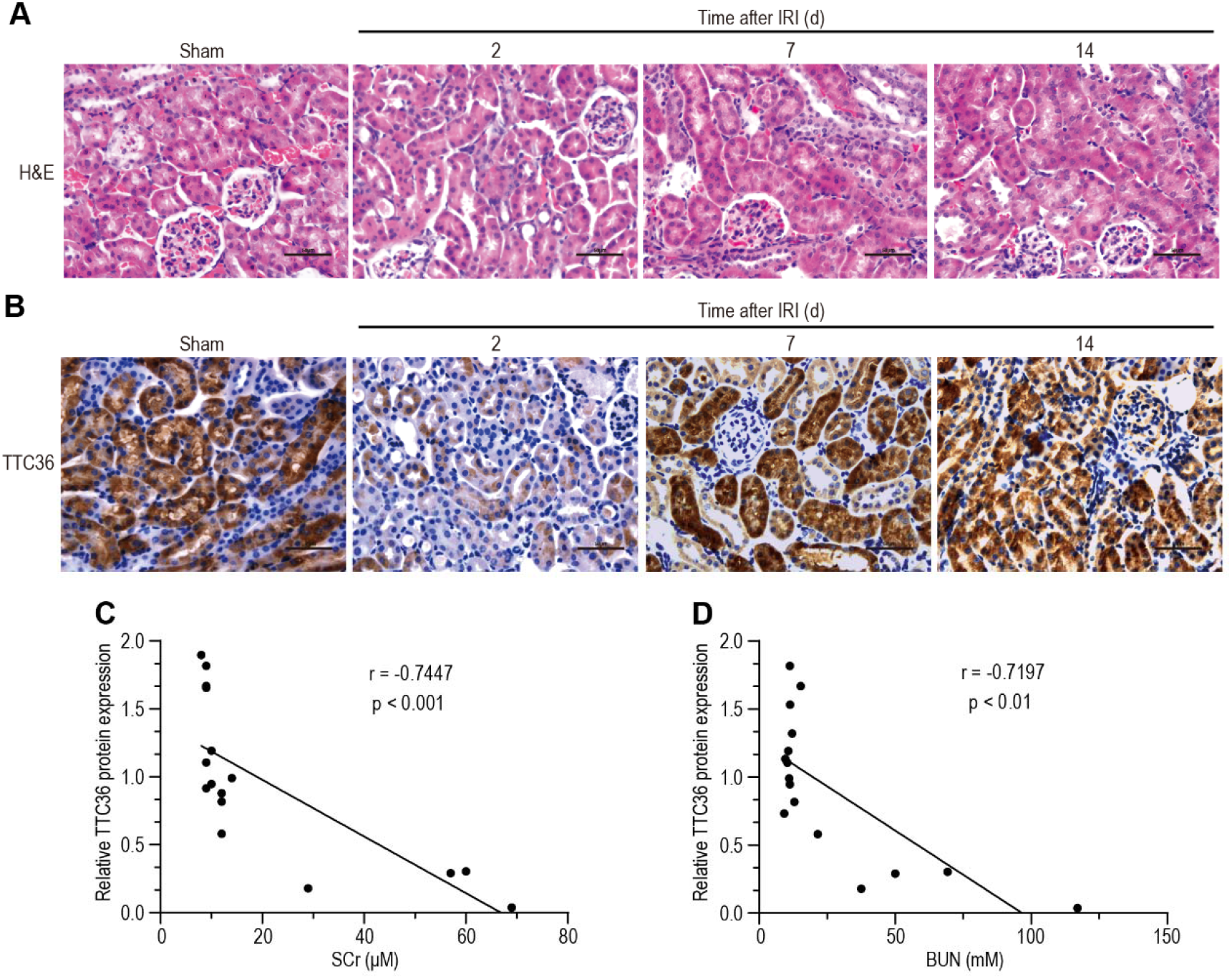
The Expression of TTC36 Was Negatively Corelated with The Degree of Kidney Injury. (A) Representative images of hematoxylin-eosin staining in the kidneys of IR-treated mice (Sham and 2, 7 and 14 days after IR-treatment; n = 3 mice per group; scale bars, 50 μm). (B) Representative images of immunohistochemical staining of TTC36 in the kidneys of IR-treated mice (Sham and 2, 7 and 14 days after IR-treatment; n = 3 mice per group; scale bars, 50 μm). (C) Correlation between renal TTC36 expression and the concentration of SCr in IR-induced mice (Sham and 2, 7 and 14 days after IR-treatment; n=4 mice per group). (D) Correlation between renal TTC36 expression and the concentration of BUN in IR-induced mice (Sham and 2, 7 and 14 days after IR-treatment; n=4 mice per group). Statistically significant differences were determined by Student’s t-test and one-way ANOVA. SCr, serum creatinine; BUN, blood urea nitrogen; IR, ischemia/reperfusion; ANOVA, analysis of variance. H&E, Hematoxylin-Eosin.

### Overexpression of TTC36 protected renal tubular cells against cisplatin-induced apoptosis

To further elucidate the role of TTC36 in AKI, proximal tubular cells (HK2) were treated with cisplatin as an in vitro model of acute tubular injury. Consistent with in vivo results, overexpression of TTC36 augmented the viability and survival of HK2 cells which were subjected to cisplatin treatment (Fig. 3, A, B, and C). The expression of BCL-2, an integral outer mitochondrial membrane protein that inhibits the cellular apoptotic death, was increased, and the expression of BAX, which functions as an apoptotic activator, was decreased in TTC36 overexpressed HK2cells (Fig. 3, D and E). Consistently, in primary tubular cells isolated from *Ttc36*^-/-^ mice kidneys, the expression of BCL-2 was down-regulated whereas the expression of BAX was up-regulated in comparison to those derived from WT mice (Fig. 3F). Furthermore, the overexpression of TTC36 down-regulated cleaved Caspase-9 and cleaved Caspase-3 expression and reduced the increment of those in cisplatin-induced HK2 cells, compared to control group (Fig. 3G). Coincident with the above data, an evident up-regulation of cleaved Caspase-9 expression was observed after TTC36 was silenced (Fig. 3H). In conclusion, these results demonstrated that TTC36 overexpression was beneficial to renal tubular cells against cisplatin-induced apoptosis.

**Figure 3.**
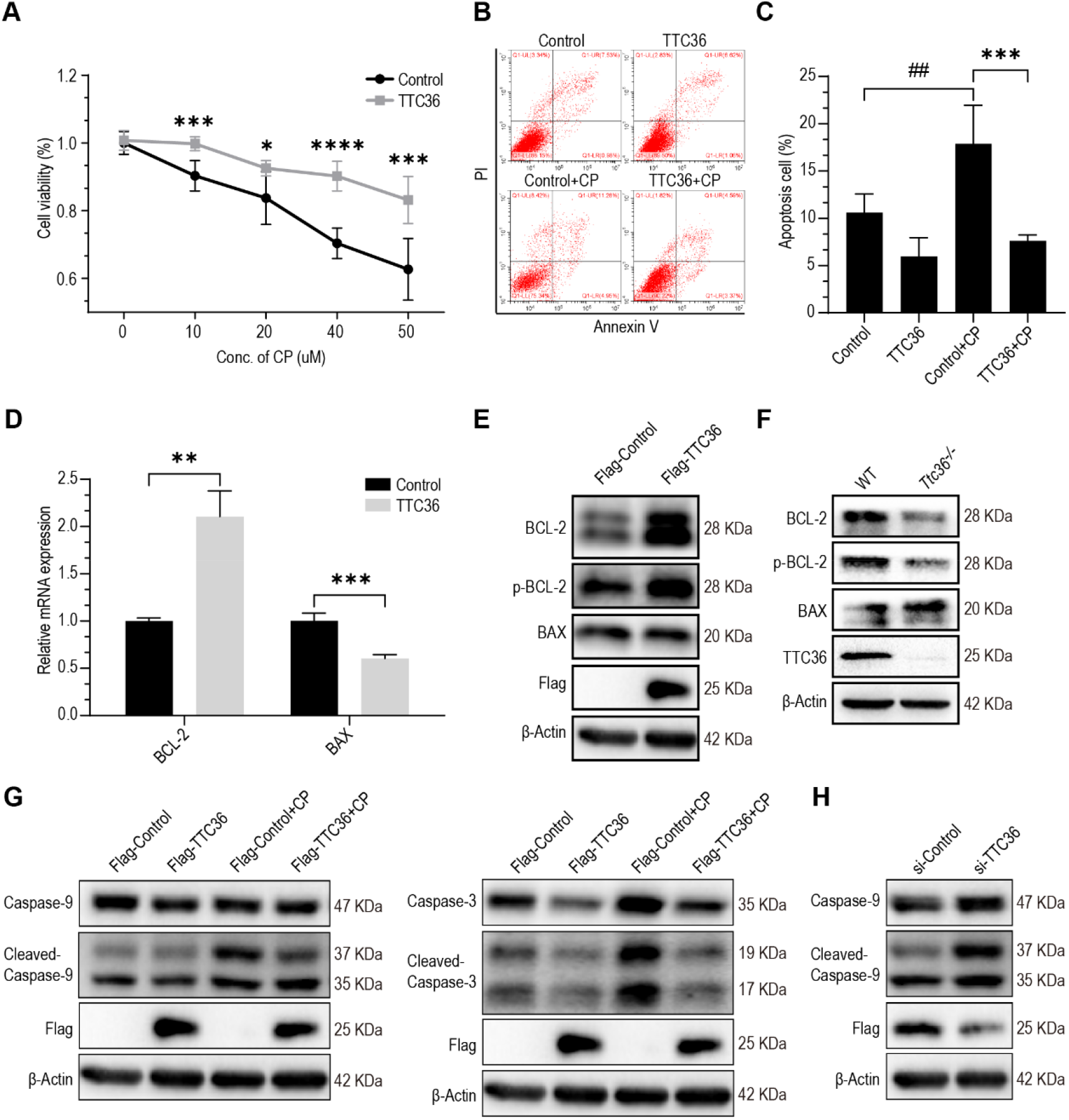
TTC36 overexpressing protected renal tubular cells against cisplatin-induced apoptosis. (A) The viability of HK2 cells with or without TTC36 overexpression were assayed with MTT after they were treated with indicated concentration of CP for 20 hours. (B) Representative images for FACS analysis after annexin V and PI staining. TTC36 was overexpressed in HK2 cells followed by treatment with 25 μM CP for 20 hours. (C) Quantitative analysis for the percentages of apoptosis cells with FACS. (D) The relative mRNA expression of *BCL-2* and *BAX* were analyzed using RT-qPCR. 18s was used as an internal control. (E) Western blotting for Flag-tagged TTC36, BAX, BCL2, and p-BCL2 in HK2 cells overexpressed with TTC36, β-Actin as a loading control. Bcl2 was detected following p-Bcl2 being stripped with stripping buffer. (F) Western blotting for TTC36, BAX, BCL2, and p-BCL2 in isolated primary tubular cells of WT and *Ttc36*^-/-^ mice. Bcl2 was detected after p-Bcl2 being stripped. (G) Western blotting for caspase-9, cleaved caspase-9, casepase-3, cleaved caspase-3, and Flag-tagged TTC36 in CP-treated HK2 cells with or without TTC36 overexpression. Caspase-9 and casepase-3 were detected in two gels. (H) Western blotting for Flag-tagged TTC36, caspase-9 and cleaved caspase-9 in CP-treated overexpressed with TTC36 HK2 cells with or without TTC36 silenced. Data are shown as means + SD (n = 3). Statistically significant differences were determined by Student’s *t*-test and one-way ANOVA. \**p* < 0.05, \*\**p* < 0.01, \*\*\**p* < 0.001, and \*\*\*\**p* < 0.0001. Results are representative of at least three independent experiments. CP, cisplatin; TTC36, tetratricopeptide repeat domain 36; BCL-2, BCL2 apoptosis regulator; BAX, BCL2 associated X, apoptosis regulator; p-BCL2, phosphorylated-BCL2 apoptosis regulator; MTT, methyl thiazolyl tetrazolium; FACS, fluorescence-activated cell sorting; PI, propidium iodide; RT-qPCR, quantitative real-time PCR; ANOVA, analysis of variance.

### TTC36 mitigated cisplatin-induced mitochondrial dysfunction

The homeostasis of mitochondria is critical for the survival and function of renal tubular cells in AKI. We further investigated the effect of TTC36 on mitochondrial membrane potential (MMP) maintenance in cisplatin-induced tubular cell injury using JC-1, which is a MMP indicator that forms aggregate and emits red fluorescence in relatively high MMP. As shown in Fig. 4, A and B, the MMP was reduced in cisplatin-treated cells and partially reversed by the overexpression of *TTC36*. Mitophagy and proper mitochondrial dynamics are vital for normal mitochondrial function. The expression of *MFN1, MFN2*, and *OPA1*, which indicate the fusion of mitochondria(18,19), was rescued by TTC36 overexpressing in cisplatin-treated HK2 cells, compared to the CP-treated control group (Fig. 4C). Based on the importance of mitophagy clearance for mitochondrial homeostasis, we examined expression of mitophagy-related genes in cisplatin-treated HK2 cells and observed that overexpression of *TTC36* alleviated the reduction of *ATG5, ATG7, PARKN*, and *BNIP3L* expression (Fig. 4D), which are responsible for the regulation of mitophagy(20–23). The decreased expression of *PPARGC1A*, *SOD2*, and *NFE2L2*, which play a protective role in mitochondria(24–27) in cisplatin-treated HK2 cells, was ameliorated by TTC36 overexpressing (Fig. 4E). Collectively, these results suggested that TTC36 plays a protective role in cisplatin-induced mitochondrial dysfunction via maintaining mitophagy clearance and mitochondrial dynamics.

**Figure 4.**
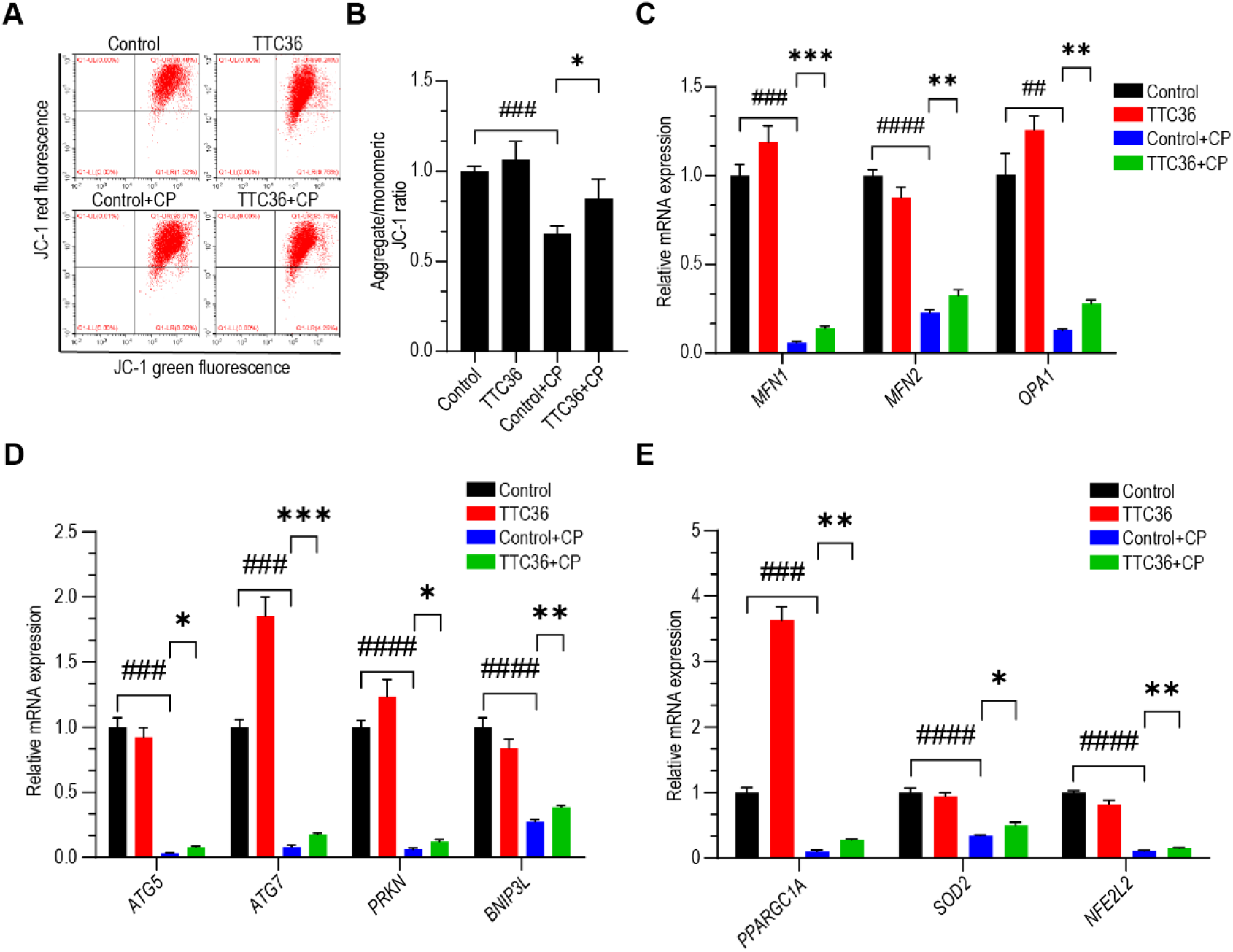
TTC36 mitigated cisplatin-induced mitochondrial dysfunction. (A) Representative images for FACS analysis after HK2 cells were incubated with CP (25 μM) for 20 hours followed by JC-1. (B) The ratios of JC-1 red fluorescence to JC-1 green fluorescence in CP-induced HK2 cells were quantified using FACS. (C) RT-qPCR analyses for *MFN1*, *MFN2*, and *OPA1* mRNA expressions. (D) The relative mRNA expressions of *ATG5, ATG7, PRKN*, and *BNIP3L* were analyzed by RT-qPCR. (E) RT-qPCR analyses for *PPARGC1A, SOD2*, and *NFE2L2* mRNA expressions. Data are shown as means + SD (n = 3). Statistically significant differences were determined by Student’s *t*-test and one-way ANOVA. \**p* < 0.05, \*\**p* < 0.01, \*\*\**p* < 0.001, and \*\*\*\**p* < 0.0001. Results are representative of at least three independent experiments. *MFN1*, mitofusin 1; *MFN2*, mitofusin 2; *OPA1*, OPA1 mitochondrial dynamin like GTPase; CP, cisplatin; *ATG5*, autophagy related 5; *ATG7*, autophagy related 7; *PRKN*, parkin RBR E3 ubiquitin protein ligase; *BNIP3L*, BCL2 interacting protein 3 like; *PPARGC1A*, PPARG coactivator 1 alpha; *SOD2*, superoxide dismutase 2; *NFE2L2*, nuclear factor, erythroid 2 like 2; FACS, fluorescence-activated cell sorting; CP, cisplatin; RT-qPCR, quantitative real-time PCR; ANOVA, analysis of variance.

## Discussion

It has been demonstrated that the specific expression of TTC36 in renal proximal tubular cells in our previous work(15). However, no study has been performed to uncover the role of TTC36 in AKI. We conducted these experiments with genetic and pharmacological methods to revealed the role TTC36 played during the pathogenesis of acute renal damage. Here, we demonstrated the pathogenic impact of TTC36 deficiency in acute renal tubular damage and mitochondrial disorder in vitro.

Overexpression of TTC36 effectively mitigated mitochondrial disorder and inhibited cell apoptosis induced by cisplatin. All these experimental data suggested the function of TTC36 in maintaining mitochondrial homeostasis, facilitating the protection for kidneys in acute injury.

It is considered that the main victims in AKI, including cisplatin-induced nephrotoxicity, are renal tubular cells, which are full of mitochondria(28). The latter are anticipated to be the primary objects in acute tubular cell injury, featured with the reduction of MMP and OCR and the impairment of mitophagy and fatty acid-oxidation(10). In consistence with the above opinion, many studies focusing on maintaining the homeostasis of mitochondrial to improve AKI have been performed(29–31). Here, we revealed that the indicators related to mitochondrial function and homeostasis, containing the MMP, the expression of mitochondria-related genes, and mitophagy were partially promoted by TTC36 overexpression in cisplatin-treated HK2 cells, suggesting that TTC36 mitigate cisplatin-induced mitochondria dysfunction.

As a pathological phenomenon and a pathogenic factor, Mitochondrial dysfunction contributes to a serious of harmful reactions, including oxidative stress, inflammation, and tubular cell damage and apoptosis(32,33). Mitochondria, which are in charge of producing a dominant portion of cellular energy in the shape of ATP, are considered as vital organelles of eukaryotic cells, including ROS level regulation, buffering cytosolic calcium, and apoptosis regulation, and those are closely related to the pathophysiology of diseases(34). Mitochondria contain bilayer membranes where the inner membrane in charge of mitochondrial oxidative phosphorylation and the outer membrane consisting of crucial proteins associated with the regulation of apoptosis. We observed that TTC36 protected HK2 cells against apoptosis in an in vitro model of cisplatin-induced acute tubular injury. However, to reveal the detailed mechanism of TTC36 in regulating cell apoptosis, there are still lots of attempts need to perform in the future.

The limitation of our research is that the role of TTC36 in apoptosis and mitochondrial homeostasis was observed in cultured cells instead of human tissue. We revealed that the down-regulation of TTC36 is negatively correlated with the degree of renal injury in vivo model of IR-induced AKI, however, the usefulness and significance of TTC36 in protecting AKI patients still need further investigations using patient specimens and clinical trials of TTC36 agonist in the future.

## Conclusions

In this study, we first discovered that TTC36 overexpression protected HK2 cells against cisplatin-induced apoptosis via maintaining mitochondrial homeostasis in the aspect of mitochondrial membrane potential and mitophagy-related gene expression. These findings uncovered in our study not only augment our understanding of the molecular mechanism and pathogenesis of AKI but also imply that developing safe and effective agonists of TTC36 could be a potential therapeutic strategy for AKI patients.

## Supplementary Material

Supplementary Table S1

